# Characterizing the relation of functional and Early Folding Residues in protein structures using the example of aminoacyl-tRNA synthetases

**DOI:** 10.1101/290627

**Authors:** Sebastian Bittrich, Michael Schroeder, Dirk Labudde

## Abstract

Proteins are chains of amino acids which adopt a three-dimensional structure and are then able to catalyze chemical reactions or propagate signals in organisms. Without external influence, most proteins fold into their correct structure, and a small number of Early Folding Residues (EFR) have previously been shown to initiate the formation of secondary structure elements and guide their respective assembly.

A dataset of 30 proteins and 3,337 residues provided by the Start2Fold database was analyzed. Proteins were represented as residue graphs in order to analyze topological descriptors of EFR. These residues constitute crucial connectors of protein regions which are distant at sequence level. Especially, these residues exhibit a high number of non-covalent residue-residue contacts such as hydrogen bonds and hydrophobic interactions. This tendency also manifests as energetically stable local regions in a knowledge-based potential. These distinct characteristics can give insights into what drives certain residues to initiate and guide the folding process. Furthermore, these features are not only characteristic for EFR but also differ significantly with respect to functional residues such as active or ligand binding sites. This unveils a split between structurally and functionally relevant residues in proteins which can improve their evolvability and robustness.

Aminoacyl-tRNA synthetases demonstrate this separation in an evolutionary context: the positions of EFR are preserved over the course of evolution and evolutionary pressure is smaller in comparison to positions relevant for protein function. The shown separation between functional and EFR has implications for the prediction of mutation effects as well as protein design and can provide insights into the evolution of proteins.

## Introduction

Most proteins adopt their three-dimensional conformation autonomously during the process of protein folding [1,2]. Various diseases are caused by misfolding or aggregation of proteins [3–6]. During the protein folding process, the denatured chain of amino acids passes an energetic barrier, called transition state, to form a compact and functional structure [2].

How proteins fold is an open question [1]. There is a lack of experimental data describing which events or residues guide the folding process [7–9]. The protein sequence resembles the starting point and the three-dimensional structure captures the result of the protein folding process for a wide range of proteins, yet how they connect via transition states is unclear. The unstable nature of transition states hinders their experimental determination [10,11]. Another obstacle for the understanding of the sequence-structure relation is that some proteins depend on chaperons to fold correctly [6].

### The defined-pathway model

Alternative folding pathways have been described for homologous proteins [12]. It is an open question if a general folding pattern can be derived which is relevant for all proteins [13]. Also, there is dispute which aspects of protein folding are stochastic and which are deterministic [14,15]. The defined-pathway model proposes that small fragments fold first and then guide a step-wise assembly of further parts of the protein until the native structure is formed [14,16,17]. Such fragments fold autonomously – no other region of the protein directly supports or hinders their formation [14,17]. Which parts of the protein initiate the formation of local, ordered structures, e.g. secondary structure elements, is encoded in their sequence [18–23]. Consequently, these regions decrease in free energy as well as entropy and stabilize the protein during the folding process [23,24]. This also supports the observation that proteins fold cotranslationally as they are being synthesized by a ribosome and stabilizing long-range contacts cannot be formed yet [25]. These local structures form long-range contacts and assemble the global structure [14,18,22,26,27]. The formation of a native structure causes a further decrease in free energy [17,28,29]. Long-range contacts are especially important for the stability of the hydrophobic core of the native structure [30].

### Identifying Early Folding Residues during protein folding

In recent years, various experimental strategies [31–34] were established which can identify residues crucial for the folding process. Pulse labeling hydrogen-deuterium exchange (HDX) [14,30,35–40] tracks the protein folding process with spatial and temporal resolution. The state of a protein can be controlled e.g. by denaturants or temperature [36]. Starting from a denatured protein, folding conditions are gradually established until the protein refolded completely. The resulting folding trajectory can be studied by HDX. Depending on the state of the folding process, individual amino acids will be susceptible to or protected from an exchange of the hydrogen atom of their amide group. Residues become protected when their amide group is isolated from the solvent as the effect of other residues surrounding them. When the folding process affects a residues, its spatial neighborhood is altered. Thereby, especially the formation of hydrogen bonds involving the amide group is relevant. Where and when these exchanges occur is tracked by a downstream mass spectroscopy or nuclear magnetic resonance spectroscopy. Residues which are protected from the exchange at the earliest stages [14,38–40] are called Early Folding Residues (EFR). Residues which are protected only at later stages or not at all are referred to as Late Folding Residues (LFR). One can also argue that the experimental signal of EFR is currently too little understood. The protection of amide groups occur at an exceedingly fast timescale. In some cases, they may not be the effect of the formation of hydrogen bonds but rather be the mere result of undirected physical chemistry. Also, other experimental techniques for the determination of key residues in the folding process [31–34] show little correlation with the annotation of EFR [23]. E.g., data from *ϕ*-value analysis is difficult to interpret on its own [31] and may differ drastically depending on the introduced amino acid substitution, so no one-to-one relation between it and EFR can be expected [30] which pronounces the difficulties of studying the structural role of EFR. They were shown to initiate the folding process and the formation of secondary structure elements [40] or even larger autonomously folding units [14]. EFR tend to be conserved, but non-functional residues [41]. In contrast, LFR may be relevant during later stages of the folding process, implement protein function, or be mere spacers between protein regions.

The data obtained by HDX experiments is difficult to interpret [42] and results of other experiments or techniques are tedious to compare [30,40]. The Start2Fold database [40] provides an invaluable annotation of EFR in a standardized manner [30]. In a previous study [39], EFR have been shown to exhibit lower disorder scores and higher backbone rigidity. Regions with relatively high backbone rigidity are likely to constitute ordered secondary structure elements and this tendency is manifested in local sequence fragments [19,20,39,40,43]. Especially aromatic and hydrophobic amino acids were linked to ordered regions of proteins [39]. Subsequently, it was shown that EFR are likely buried according their relative accessible surface area (RASA) and proposed that they are also the residues which form the greatest number of contacts in a structure [40]. EFoldMine [9] is a classifier that predicts EFR from sequence. Due to the nature of the trained models [9,39], it is still unclear what characteristics cause EFR to fold first [23]. Furthermore, early folding events are enigmatic [15,44,45]. EFR are a resource to address this question: are the experimental signals of EFR transient implying that EFR are only relevant in the early stages of the folding process or will EFR also exhibit distinct characteristics in the successfully folded, native conformation?

### Representing proteins by Energy Profiling and residue graphs

The free energy of a native protein structure is minimal [14]. Thus, knowledge-based potentials are a potent tool to describe the process of protein folding [28] and have been previously employed for the quality assessment of protein structures [29]. The complex interactions of a residue in the three-dimensional structure are expressed as single computed energy. Each amino acid has a propensity to be exposed to the solvent or be buried in the core of a protein which can be expressed as pseudo-energy according to the inverse Boltzmann law. The energy of a residue is calculated by summing up the pseudo-energy of all residues in spatial proximity (i.e. distance less than 8 Å) [28]. Low computed energies occur for hydrophobic amino acids which are stabilized by many contacts. Thus, this approach is a valuable feature to assess the stability of individual residues as well as their interactions with their spatial neighborhood.

Individual residues can also be characterized in the context of protein structures by topological features derived from network analysis. Protein structures are represented as graphs: amino acids constitute the nodes and contacts between residues are represented as edges [11,46–52]. There is a plethora of contact definitions and most are based on distance cutoffs between certain atoms of amino acids [53]. Graph representations of proteins were previously employed to describe residue flexibility [54] as well as residue fluctuation [46], protein folding [11,49], structural motifs [55], and evolvability [52]. Furthermore, residue graphs were shown to exhibit the character of small world networks [11,46–49] whereby a small number of residues has high connectivity and the average path length in the graph is small. Hydrophilic and aromatic amino acids were found to be crucial connectors in the graph – so-called hubs – which underlines their importance in the context of protein folding [56].

Graph representations of proteins also allow to assess whether proteins feature a modular design [57,58]. Similar problems are solved by similar strategies and existing, established, and safe strategies seem to reemerge [59]. This may explain why the explored sequence and structure space is relatively small: by evolving established sequences, misfolding sequences or those prone to unfavorable aggregation [60] are avoided [59,61]. This behavior is likely the result of a separation of residues relevant for folding and those relevant for protein function [41] such as ligand binding sites or active sites. Functional residues also were shown to exhibit distinct topological features [48]. This separation increases robustness and evolvability of protein sequences [41,52,57,58,62] because functional residues can be changed without any impairment of the protein’s stability and the fold can be improved without compromising function.

The Start2Fold database [40] constitutes a dataset of EFR [9,14,17,23]. Previous studies considered a small number of proteins, whereas the 30 proteins of the Start2Fold database [40] allow a more robust characterization of EFR. Because the annotation of EFR is standardized, a workflow can be established to analyze also future results of HDX experiments added to the database.

## Motivation

It is unknown what sequence features causes particular residues to fold early and how these residues contribute to the formation of the native structure (Fig 1A). EFR are connected to the defined-pathway model and provide an opportunity to understand the driving forces behind the assembly of stabilizing local structures as well as the formation of tertiary contacts [14,23].

**Fig 1.**
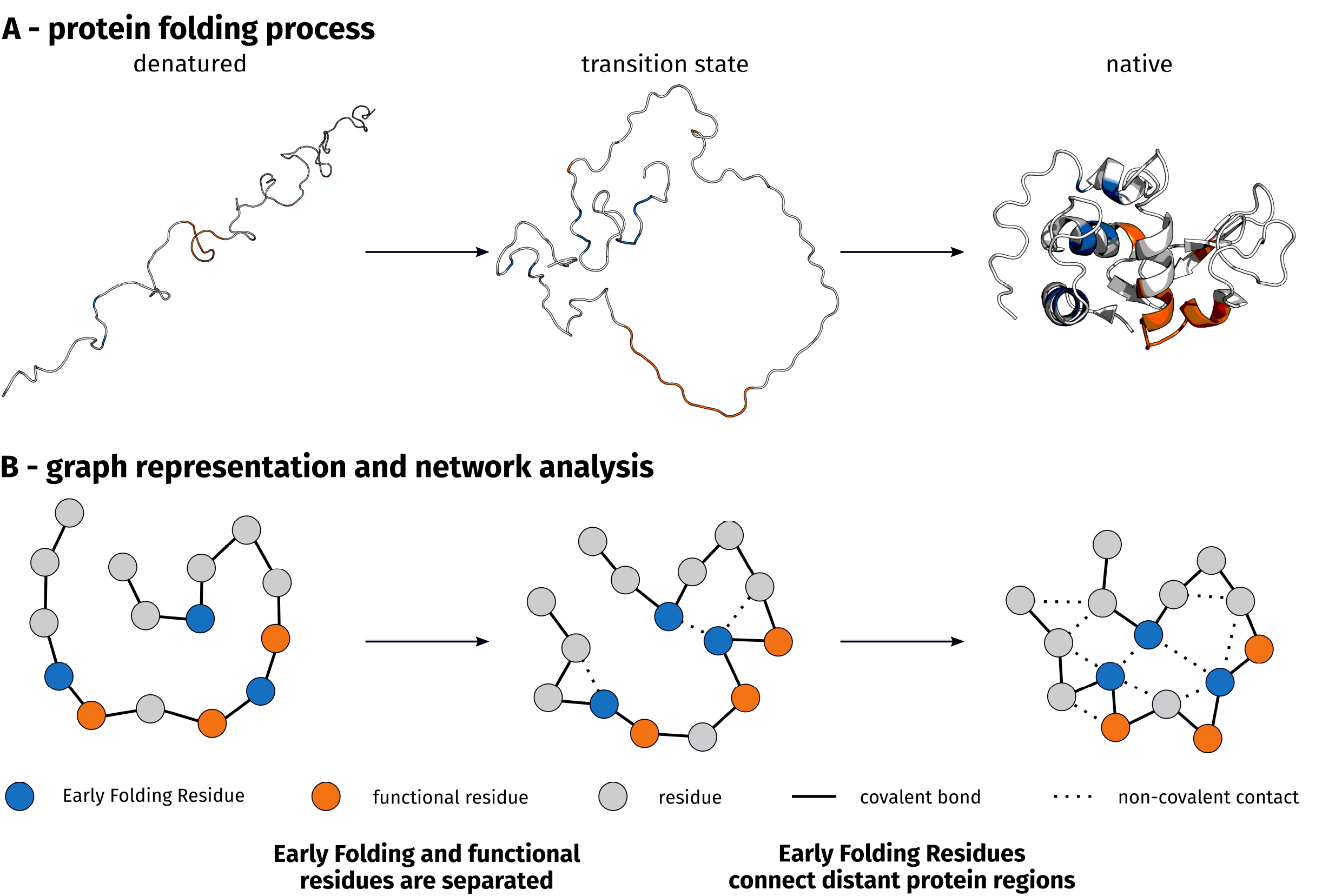
Graphical abstract. (**A**) During the folding process, an extended protein chain passes the transition state and forms a native structure [2]. (**B**) Protein structures are represented as graphs to derive topological descriptors of residues. Amino acids constitute nodes, whereas residue contacts are represented as edges. EFR are structurally relevant residues which participate early in the folding process by forming local contacts to other residues. They are separated from functional residues which are primarily ligand binding sites and active sites as derived from UniProt [63]. EFR show a great number of long-range contacts which furnish the spatial arrangement of protein parts which are far apart at sequence level.

In this study, several novel structural features are employed for the characterization of EFR. Especially, the Energy Profiling approach, topological descriptors of residue graphs, and the explicit consideration of non-covalent contacts types provides a new level of information in order to describe the folding process. EFR exhibit lower, more stable computed energies in their Energy Profile [28,29]. Network analysis reveals that EFR are more connected to other residues and that they are located at crucial positions in the residue graph (Fig 1B). This distinct wiring to the rest of the protein is especially furnished by hydrophobic interactions. EFR are likely structurally relevant for the correct protein fold [9]. This information is used to demonstrate that proteins separate structurally relevant residues from functional residues (Fig 1B). We show that positions of EFR are preserved over the course of evolution in the diverse superfamily of aminoacyl-tRNA synthetases (aaRS). Again a separation between EFR and functional residues is evident. Furthermore, positions of EFR are less sequentially conserved than positions relevant for protein function. Interestingly, EFR are in all cases predicted to be located in the center of secondary structure elements.

## Results and Discussion

A previously described dataset [23] of 30 proteins and 3,377 residues is the basis of this study and summarized in S1 Table. 482 (14.3%) of the residues are labeled as EFR, the remaining residues are considered LFR. Hydrophobic amino acids have a higher propensity of being EFR (S1 Fig) as previously described in literature [23].

To characterize EFR in more detail, various features were defined and compared to the values of LFR. EFR form a significantly greater number of residue-residue contacts (i.e. distance less than 8 Å) than their LFR counterparts (Fig 2A). The loop fraction is defined as the ratio of unordered secondary structure elements in a window centered on a particular residue [64]. Fewer unordered secondary structure elements can be found around EFR (Fig 2B), whereas LFR exhibit a higher propensity to occur in coil regions. EFR are on average closer to the centroid of a protein structure and are likely embedded in the hydrophobic core (Fig 2C). Analogously, they also tend to be more distant to the N- or C-terminus of the sequence than other residues and are likely buried regarding their RASA as per S2 Table.

**Fig 2.**
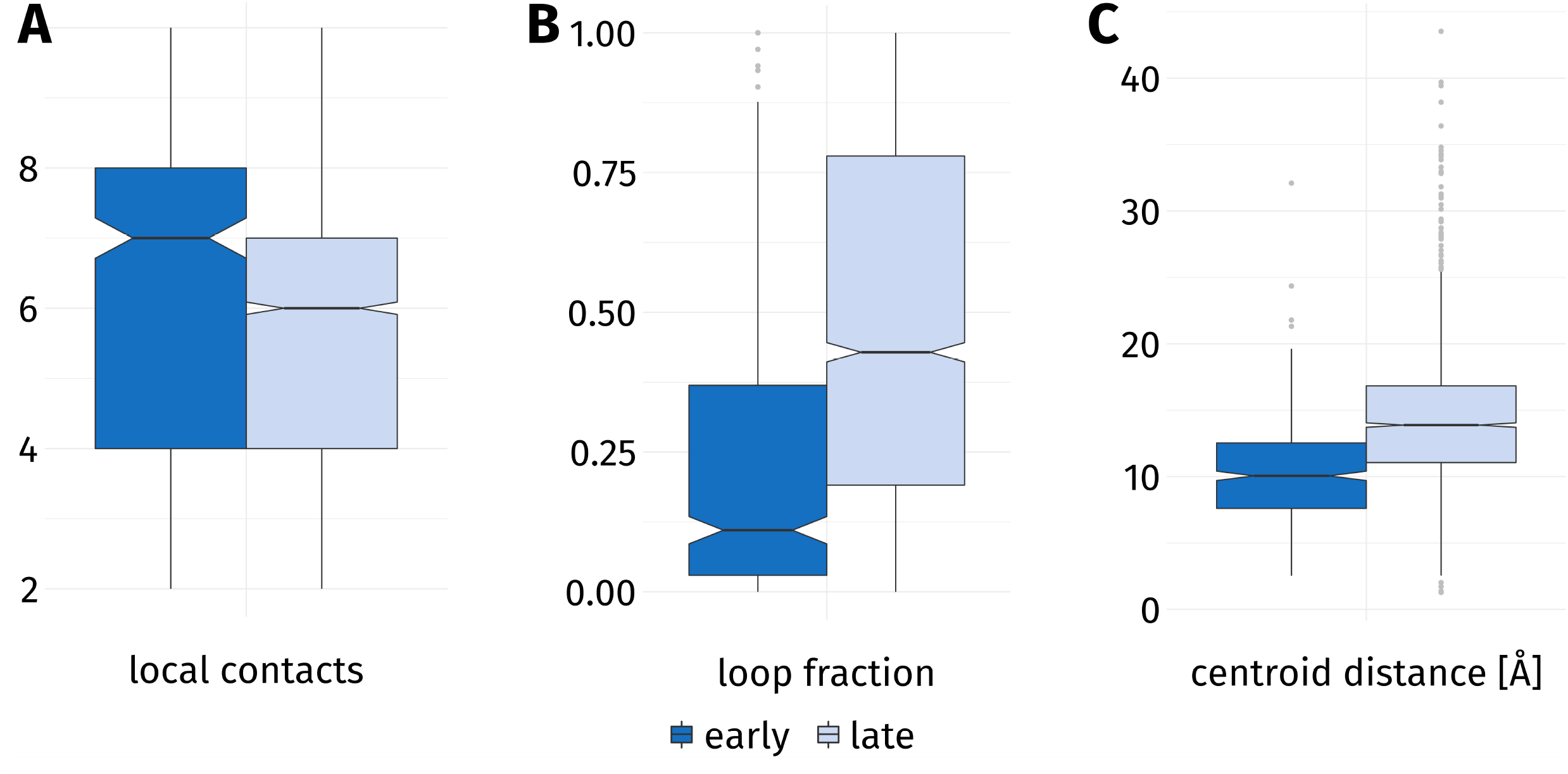
General properties of Early (dark blue) and Late Folding Residues (light blue). (**A**) EFR form more contacts to their surroundings than LFR. (**B**) The loop fraction [64] is the ratio of unordered secondary structure elements which are observed in a windows of nine amino acids around a residue. EFR are more commonly surrounded by ordered secondary structure elements. (**C**) EFR are located significantly closer to the centroid of the protein than LFR.

The propensity of EFR to participate in more contacts and to occur in the core of a protein are in agreement with previous studies [14,23,39,49]. The shift in loop fraction can also be attributed to these findings and is further substantiated by the fact that long ordered secondary structure elements tend to contain more EFR [23]. It has been reported that buried residues are more likely to be EFR [23,30] which also explains why they are closer to the spatial centroid of a protein and more separated from sequence termini (S2 Table). Evolutionary couplings scores reported by the direct couplings analysis [65,66] and evolutionary information exhibit interesting properties: the reported coupling strength as well as evolutionary information of EFR is significantly increased. The relation of evolutionary information and EFR has been the subject of previous studies [9,23]. Correlations between features are presented in S3 Fig, e.g. the contact count as well as the evolutionary coupling descriptors are strongly correlated. All these factors can neither explain why some residues become EFR while others do not nor how EFR relate to the rest of a protein in terms of network analysis.

### Early Folding Residues constitute stable local conformations

To assess the energetic contribution of EFR to the native structure, the proteins of the dataset were transformed by the Energy Profiling approach [28,29]. The computed energy of EFR are significantly lower than the values of LFR. A more detailed investigation of computed energy (Fig 3 and S2 Fig) shows that this trend can be observed for individual amino acids, but the change is insignificant for aspartic acid and isoleucine. Hydrophilic amino acids commonly feature high computed energies, whereas the values for hydrophobic and aromatic amino acids are low. The changes in computed energy for amino acids with hydroxyl groups in their side chain such as serine and threonine are remarkable. Futhermore, cysteine stands out with a high variance of computed energy in the LFR state and low variance for EFR. Energy values predicted from sequence using the eGOR method [28] are also lower for these residues (see S2 Table) which indicates that the position of EFR is the consequence of the sequence composition of small fragments. Regarding the average absolute contact frequencies, a EFR participates in 3.87 hydrogen bonds and forms 1.30 hydrophobic interactions with other residues. This constitutes a significant increase compared to LFR (see S2 Table).

**Fig 3.**
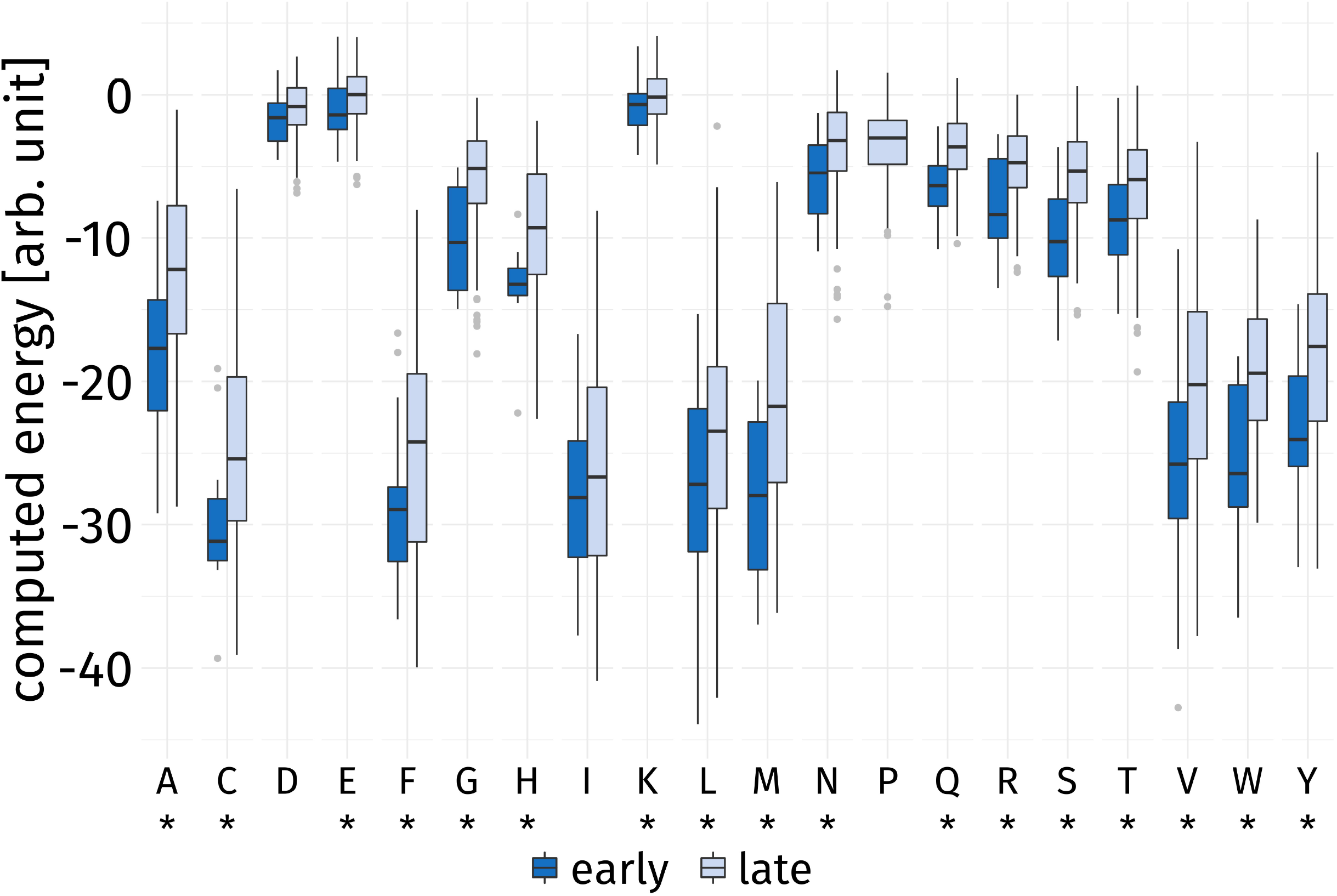
Computed energy by amino acid. The Energy Profiling approach [28,29] was used to characterize the surrounding of each residue. Hydrophobic and aromatic amino acids have a high tendency to be located in the buried core of a protein. Hydrophilic and polar amino acids prefer to be exposed to the polar solvent. This tendency is reflected by low and high average computed energies respectively. The distribution of computed energies of EFR always exhibits a lower median than LFR. Significance in change is indicated by asterisks (*). EFR observations of serine and threonine exhibit relatively low computed energies. The side chains of both amino acids can form hydrogen bonds. The decrease in computed energy is insignificant for aspartic acid and isoleucine. No annotation of EFR is available for proline.

The results indicates that EFR occur in parts of proteins which are more stable and contain an increased number of hydrophobic amino acids in their spatial surroundings. Especially amino acids such as serine or threonine, which can form hydrogen bonds via their side chains, feature relatively low computed energies even though they have an overall tendency to be exposed to the solvent due to their hydrophilic nature. The energy contribution of hydrogen bonds has been shown to be context-specific [67], but also crucial for the correct formation of protein structure [56]. Especially amino acids capable of forming side chain hydrogen bonds contribute to the protein stability [1,67]. Hydrophilic and aromatic amino acids like arginine, histidine, and methionine are considered strong hubs in protein structures [56], which is substantiated by a significant change in computed energy for EFR. Furthermore, arginine and histidine can form hydrogen bonds by their side chain as well. Hydrophobic amino acids occur in the core of a protein and are stabilized by an increased number of hydrophobic interactions. Almost all amino acids experience a significant decrease in computed energy in the EFR state which probably relates to their specific preferences being fulfilled. E.g., hydrophobic amino acids in a hydrophobic environment have favorable computed energy and the same is true for polar amino acids if they favorably interact with similar residues. A wide range of biological roles has been reported for cysteine. It is relevant for ligand binding sites as well as metal coordinating motifs [68] and is also well known for structure stabilization by disulfide bridges especially in extracellular proteins [69]. Also, a strong hydrophobic tendency has been observed in the reduced state [70], which explains why cysteines, when they are also EFR, commonly occur in the hydrophobic core and exhibit low, stable computed energies. How the hydrophobic core of a protein is established is still in debate [1,71]. There are cases where the hydrophobic collapse to a molten globule precedes the formation of secondary structure elements [72]. The low computed energies indicate that EFR have an intrinsic propensity to form stable, local conformations. EFR might be the mediators between the formation of local structure elements and their assembly in the context of the three-dimensional structure. Secondary structure elements such as helices interact e.g. by hydrophobic interactions [73], however, it seems that single contacts are neither strong nor specific enough to guide their assembly [17,74,75]. A fine-grained distinction of contact types including *π*-stacking and hydrophobic interactions would be required to assess the role of EFR as potential driving force behind the correct of arrangement of secondary structure elements.

### Network analysis shows a unique wiring of Early Folding Residues

The way residues interact with their spatial surrounding was assessed by network analysis based on residue graphs. Regarding the topological properties of residues derived from network analysis (see S4 Fig for a graphical depiction), EFR show a higher interconnectedness than LFR. They exhibit higher betweenness (Fig 4A) and closeness (Fig 4B) values. High betweenness values are observed for well-connected nodes which are passed by many of the shortest paths in a graph. High closeness values occur for nodes which can be reached by relatively short paths from arbitrary nodes. The distinct neighborhood count expresses to how many sequentially separated regions of a protein a residue is connected. Again a significant increase can be observed for EFR (Fig 4C). Residues are considered separated when they are more than five sequence positions apart. This threshold was also used to distinguish local contacts (i.e. less than six residues apart) and long-range contacts. Interestingly, the clustering coefficient features a significant decrease when EFR are considered. The clustering coefficient of a node is the number of edges between its adjacent nodes divided by the theoretical maximum of edges these nodes could form. However, EFR are biased to be in the core of the protein [40], thus, it was assessed if this change is also significant when only buried [76] residues are considered. The differences are insignificant in that case (see S2 Table).

**Fig 4.**
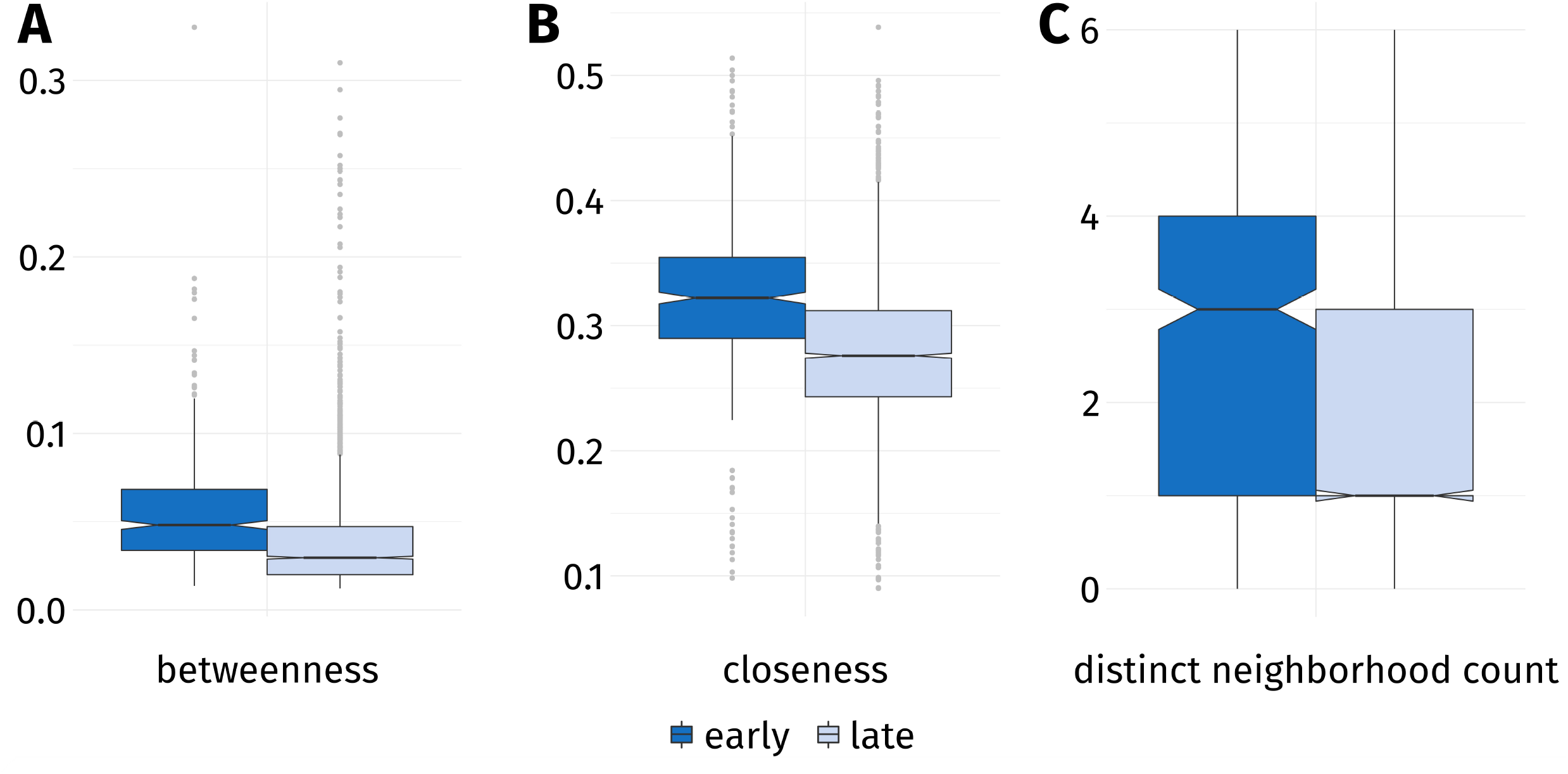
Topological properties of EFR and LFR. Proteins were represented as residue graphs and a network analysis was performed. (A) EFR have higher betweenness values implying that shortest paths in the graph tend to pass through these nodes more often. (B) They also exhibit higher closeness values because their average path length to other nodes is lower on average. (C) The distinct neighborhood count of a residues describes to how many separated regions it is connected. Residues are considered separated when their separation at sequence level is greater than five. EFR connect significantly more regions of a protein than LFR.

The betweenness property is closely related to the small-world characteristics of networks (i.e. they are well-connected even when between most nodes no edge is present) and can be observed in this case due to the ratio of protein surface and volume [49]. Residues relevant for the folding process have been shown to exhibit high betweenness values in the transition state and to be crucial for the formation of the folding nucleus [49]. Interestingly, the clustering coefficient shows no difference between EFR and LFR when only buried residues are considered. Also, the value is higher for LFR, which is probably an effect of EFR being hubs which connect several separated regions of a protein (as shown by the distinct neighborhood count). These regions themselves are not well-connected, which results in a lower clustering coefficient for EFR. The performed network analysis aids the understanding on the idiosyncratic properties of EFR in the context of the whole protein and is in agreement with previous studies [10,49,56]. EFR are hubs between sequentially distant protein regions which underlines their importance for the correct assembly of the tertiary structure of a protein. Nevertheless, the increased number of local and long-range contacts of EFR point to their importance for the whole protein folding process as described by the defined-pathway model [14,17]. The protein folding process is difficult to study due to various aspects such the existence of disordered proteins [6,39], the relevance of chaperons [6], cotranslational folding [25], and the insertion of membrane proteins by the translocon [73,77]. EFR are a welcome simplification to advance the understanding of the protein folding process.

### Early Folding Residues are non-functional residues

Division of labor is one of the most successful strategies of evolution [41,57,58,78–81]. The separation of residues crucial for folding and those furnishing function may allow reuse of established protein folds [33,41,57–59,62]. The sequence and structure space ascertained over the course of evolutions seems small for a truly random exploration. Reusing established folds could also avoid slow-folding sequences or those prone to aggregation [32,44,59,82]. There seem to be a delicate balance in proteins between robustness and evolvability [58,62,71]. Functional residues [83] can be mutated and new functions can be adopted without compromising the fold of the protein [33]. In consequence, a clear division should be observable between EFR – which initiate and guide the folding process - and the functional ones implementing protein function.

To address this question, residues in the dataset were labeled as either EFR or LFR as well as being either functional or non-functional. Active sites and ligand binding regions were considered to be the functional parts of proteins. The distribution of both binary variables (Table 1) shows that the majority of residues in the dataset are neither EFR (87.2%) nor functional (95.4%) residues. Only 0.5% share both classes, whereas 0.8% are expected to share both classes if their association was random (see methods for details). The distribution of both variables separated by individual proteins is presented in S1 Table. For most proteins, no residues are both EFR and functional (Fig 5A). Furthermore, EFR tend to be located in the core of proteins, whereas functional residues are exposed towards the solvent in order to realize their respective function (Fig 5). The acyl-coenzyme A binding protein (STF0001) [34,84,85] features five residues which are both EFR and functional (Fig 5B).

**Table 1.** Contingency table of folding characteristics and functional relevance.

|  | functional | non-functional |
| --- | --- | --- |
| early | 14 | 345 |
| late | 116 | 2332 |

Out of 2807 observations, 0.5% are EFR and functional at the same time. Based on the presented frequencies, 0.8% of all residues are expected to share both labels if their association was independent. This captures the tendency of EFR to not be functional and vice versa. Proteins were excluded when no annotation of functional residues existed.

**Fig 5.**
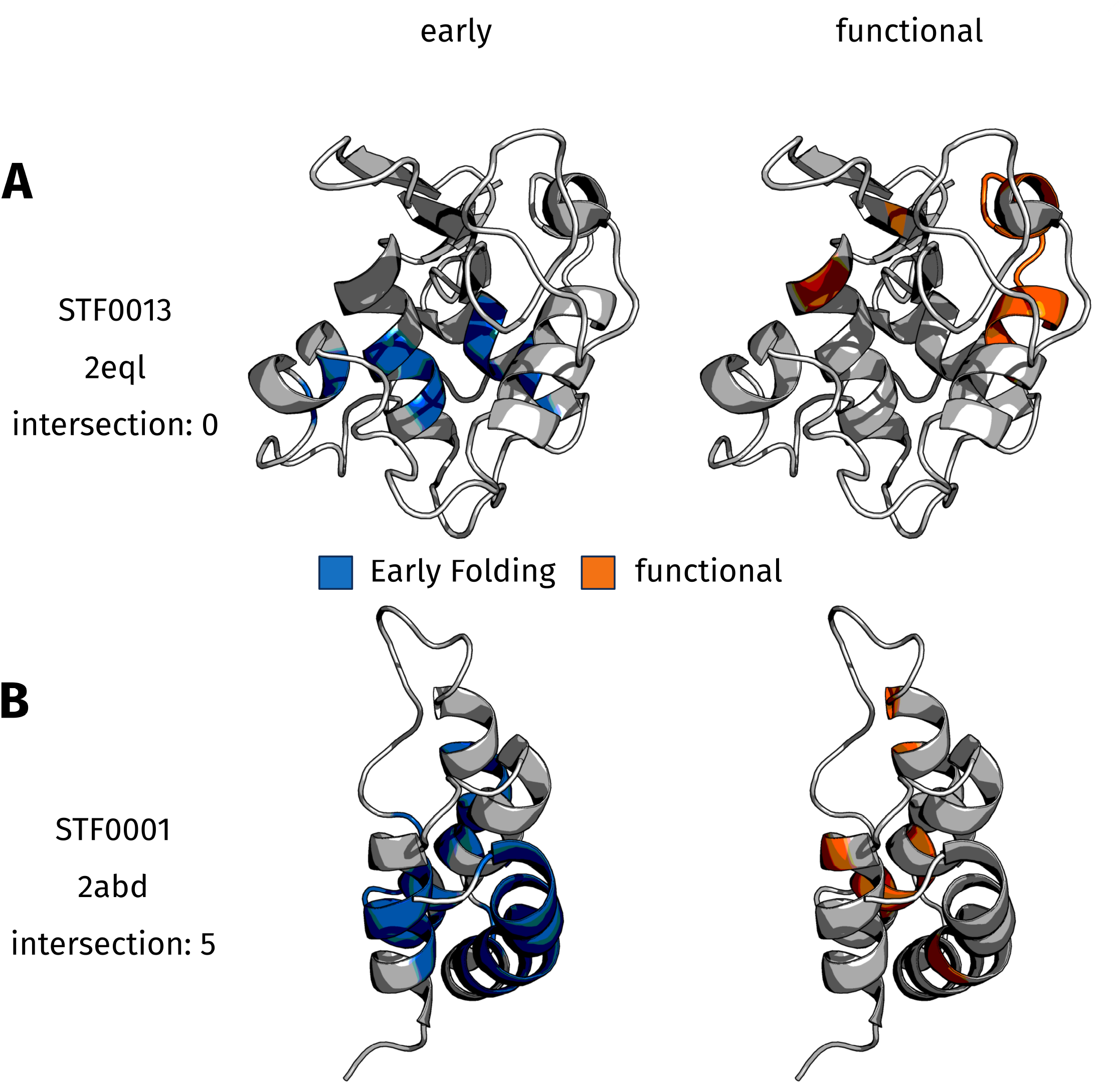
Rendered structures of 2 dataset entries. EFR are rendered in blue, functional residues are rendered in orange. (**A**) In the case of lysozyme (PDB:2eql_A) the intersection of EFR and functional residues is empty. For most proteins in the dataset, there is a clear distinction between both classes and structurally relevant residues have a propensity to be located in the core, while functional residues are exposed on the protein’s surface. (**B**) Five residues are both EFR and functional in the acyl-coenzyme A binding protein (PDB:2abd_A) which is one of the exceptions in the dataset where some residues are both EFR as well as functional.

For the majority of the dataset, a clear separation of EFR and functional residues can be observed. The acyl-coenzyme A binding protein may exhibit five residues which are both EFR and functional because its a rather small protein of 86 residues which binds ligands with large aliphatic regions. Intuitively, the residues furnishing the bowl-like shape of the protein are also those which participate in the function of ligand binding [34,84,85]. Roughly half the residues of the acyl-coenzyme A binding protein are marked as EFR which further accentuates why the separation is less strict in this case. Exceptionally well separated are EFR and functional residues in the fibroblast growth factor 1 (STF0024, S5 FigA) and Villin-1 (STF0028, S5 FigB). The first protein contains a large number of EFR distributed throughout the sequence and a large heparin-binding region which are distinct at sequence level. Villin-1 exhibits a similar distribution of EFR and features a C-terminal polyphosphoinositide binding region which contains no EFR. In both cases, the functional sites bind other molecules. This characteristic is commonly associated to increased structural flexibility [86] which may explain why EFR rarely occur there. The primary selection pressure during evolution is on protein function [87] rather than on structural integrity [88]. In cases where a certain position is crucial for function, slower folding is tolerated which implies that structure and folding are subordinated to function [71]. Disordered proteins are another example of proteins without structural integrity which achieve a high robustness of function [52]. In structural biology, structure is commonly considered to be equal to function [52,87]. However, ultimately it is most important that proteins are functional [87,89]. This potential irrelevance of a particular fold underlines that the separation of structurally and functionally relevant residues may be advantageous for evolvability. However, also cases were described where it is advantageous to place functional residues close to residues ensuring structural integrity in order to maintain protein function over the course of evolution [90]. Another interpretation with respect to the defined-pathway model [14] is that EFR initiate and guide the folding process. By assigning this responsibility to a small number of residues, the remaining residues can constitute active sites.

### Early Folding and functional residues exhibit distinct features

The previously described features were employed to substantiate the identified separation of structure and function at residue level (S3 Table). EFR show significantly lower computed energies when compared to LFR or functional residues (Fig 6A). Functional residues exhibit higher computed energies than their non-functional counterparts. Most residues form only a small number of hydrophobic interactions, however, the number for EFR is significantly increased (Fig 6B). 97.7% of EFR form hydrogen bonds and 64.3% participate in hydrophobic interactions. Functional residues participate to 93.1% in hydrogen bonds and to 43.8% in hydrophobic interactions. On the contrary, the change between the hydrogen bond count of EFR and functional residues in a buried state is insignificant (S3 Table). The clustering coefficient of a node captures how many edges can be observed between the adjacent nodes and, thus, describes how well-connected the direct surroundings of a node are. Functional residues show an insignificant change regarding this property (S3 Table). In contrast, the clustering coefficient significantly decreases when EFR are compared to LFR or functional residues (Fig 6C). In summary, EFR exhibit distinct properties compared to functional residues. Their surrounding secondary structure elements, values in Energy Profiles, and the number of hydrophobic interactions are especially characteristic. In terms of evolutionary information, functional residues exhibit a significant change compared to non-functional residues (S3 Table). When buried EFR evolutionary information of functional residues amounts to 49.11 compared to 42.04 for EFR which constitutes a significant increase. LFR and non-functional residues are less conserved at sequence level.

**Fig 6.**
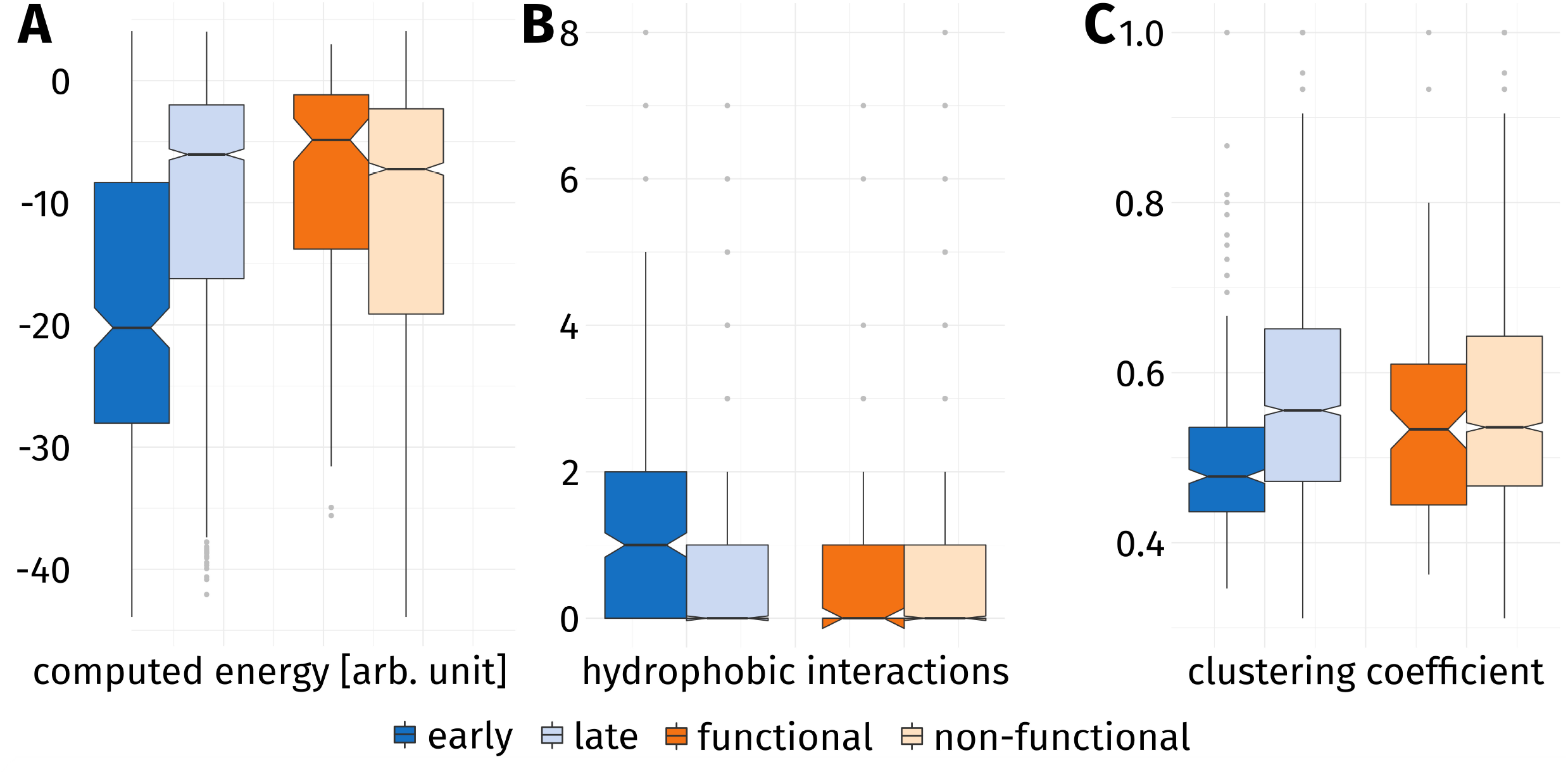
Characteristics of EFR and functional residues. EFR (dark blue) and LFR (light blue) are compared to functional (dark orange) and non-functional (light orange) residues. (**A**) EFR show lower computed energies than they are in contact with many residues and tend to be embedded in the hydrophobic core. In contrast, functional residues are exposed to the solvent in order to constitute e.g. binding sites. (**B**) Hydrophobic interactions occur especially in the core of a protein, thus, most residues do not form any. However, EFR show an significant increase compared to LFR. (**C**) The clustering coefficient of a node describes how well-connected its adjacent nodes are. EFR connect regions of a protein which are separated at sequence level and, thus, not well-connected on their own. Functional residues exhibit higher clustering coefficient indicating a more connected set of adjacent nodes.

Due to their purpose, EFR are located in the hydrophobic core and functional residues are primarily exposed to the solvent. These distinct requirements manifest in the computed energies. Furthermore, protein function can commonly be broken down to amino acids which feature hydrophilic, chemically functional groups [83]. Hydroxyl groups are a prominent examples for functional groups contributing to catalysis [83]. Thus, functional residues are likely to exhibit above average computed energies because of their higher propensity to contain hydrophilic side chains. Analogously, fewer hydrophobic amino acids constitute the functional residues of binding sites and they form fewer hydrophobic interactions. Most of the hydrophobic interactions are accumulated in the hydrophobic core of a protein [1,28,91]. EFR tend to be crucial connectors in proteins, however, their clustering coefficient is low. This can be attributed to the fact that EFR connect many distinct neighborhoods. Furthermore, functional residues feature above average closeness values: they are well-connected to other parts of the protein, even though they are unaffected by the early folding events. It was shown that functional residues have special requirements on how they are wired to the rest of a protein [48]: surrounding residues ensure the correct placement of functional residues [48,68,92], modulate their chemical properties such as p*K*_a_ values [48,83,93], or propagate signals to other parts of a protein [48]. Analogously, the evolutionary pressure on functional residues is increased compared to EFR and non-functional residues as indicated by the evolutionary information (S2 Table). In particular, catalytic activity of amino acids can be broken down to functional groups of their side chain [83]. The hydroxyl side chain of serine may be substituted by threonine or tyrosine. In contrast, contacts which stabilize protein structures can be primarily broken down to the hydrophobic or hydrophilic character of amino acids [94, 95] which allows for a wider range of tolerated mutations. Early stages of protein folding sample transient conformations [14,23] and settle for stable, local structures as indicated e.g. by the Energy Profiling approach. It has been shown that the characteristic of EFR is not directly linked to individual amino acids but rather the effect of the sequence composition of sequence fragments [9,23,39]. This may be another explanation why EFR are less conserved at sequence level than functional residues. That the folding nucleus of proteins is not necessarily sequentially conserved has been demonstrated previously [14,96,97], and makes it even more remarkable that coevolution techniques such as the direct coupling analysis perform so well for structure prediction tasks [65,66].

Modularity in proteins is also present in domains [57], secondary structure elements, and autonomous folding units of the defined-pathway model [17,27]. Particularized knowledge of EFR may improve synthetic biology and could allow the design of proteins combining existing functional domains without influencing one another negatively [2,57,58,98]. Furthermore, understanding the differences of structurally relevant residues and those implementing function could help in predicting mutation effects and provide a new level of detail by allowing whether a mutation disrupts the fold or the function of a protein [99,100].

### The position of Early Folding Residues is consistent in aminoacyl-tRNA synthetases

For the Start2Fold dataset [23] a separation of EFR and functional residues can be observed. However, no analysis of EFR in an evolutionary context is feasible due to limitations of this dataset. aaRS may be the protein superfamily with the most intriguing evolutionary history and, thus, are a prime candidate to analyze in the context of the previous findings as their emergence is well-discussed in literature [80,81,101–104]. aaRS enzymes attach amino acids to their cognate tRNA, which is subsequently recognized by its anti-codon and consumed by a ribosome. Thus, aaRS implement the genetic code and give insights into the earliest stages of life. For each amino acid, a dedicated aaRS implementation exists in each organism. E.g., AspRS attaches aspartic acid to tRNA^Asp^ in two-step reaction which involves the recognition of ATP, amino acid, and tRNA:

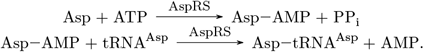

The 20 implementations can be divided into two complementary classes which differ significantly at sequence and structure level, feature distinct reaction mechanisms, and occur in diverse oligomerization states. In a recent study [104], two ligand binding motifs – the Backbone Brackets and the Arginine Tweezers – were identified, characteristic for each aaRS class. These motifs were furthermore linked to primordial implementations of both aaRS classes called protozymes [80,81]. It is hypothesized that all contemporary aaRS genes originate from the protozyme region of either class I or class II aaRS. Further analysis focuses on regions of today’s aaRS structures which correspond to the protozyme regions in order to assess how EFR predicted by EFoldMine [9] related to functional residues [104] in an evolutionary context. ATP and amino acid recognition sites were considered functional (see S6 Fig). Furthermore, we wanted to assess whether the predicted positions of EFR are consistent in these highly diverse superfamily of enzymes. This analysis is backed by a manually curated dataset which accounts for high diversity of contemporary aaRS implementations [104].

Fig 7 depicts the hypothesized protozyme [80,81,104] of each aaRS class with an aminoacyl-AMP ligand present which captures the intermediate of the enzymatic reaction. Analysis is based on 81 non-redundant structure for class I and 75 for class II, respectively. For each analyzed structure the corresponding sequence was used to predict the position of EFR [9]. A consistent numbering of residues within each class was established by a structure-guided multiple sequence alignment (MSA) [105]. Even within the depicted catalytic core of aaRS structures, sequences feature a high degree of variability and various inserts. Interestingly, residues predicted to be Early Folding are located at MSA columns which may not be extraordinarily conserved but are present in at least half of the corresponding sequences. Despite the high sequence variability of aaRS proteins, EFR positions are mostly conserved among homologues. ATP binding sites are also consistent for the structures, whereas the exact position of amino acid binding sites varies drastically. In the visualized protozyme regions (Fig 7), positions of EFR are located in ordered secondary structure elements. Functional residues, especially those realizing ATP recognition, are located in spatial proximity to one another. Furthermore, they occur in unordered coil regions and are located close to the ligand. ATP binding sites (dark orange) can be found on the left in proximity of the adenine part, whereas amino acid recognition sites (light orange) can be found on the right close to the amino acid part of the ligand. The average sequence conservation of the protozyme regions is 1.59 (1.42) for class I (and class II respectively). Positions predicted to be EFR exhibit scores of 2.50 (2.80). That for ATP binding sites is 3.75 (3.75) and for amino acid binding sites 1.85 (2.17). On average the EFoldMine prediction is 0.09 (0.09) for the protozyme regions. Positions considered EFR exhibit high values of 0.21 (0.20). ATP binding sites feature low scores, whereas amino acid binding sites feature slightly increased probabilities of being EFR (summarized in S4 Table). Detailed data for the annotated EFR and functional positions is provided in S4 File and S5 File. Because the position of amino acid binding sites is not consistent in the MSA, sequence conservation of these positions is relatively small. In contrast, ATP binding sites are mapped consistently in the MSA for both aaRS classes [104]. EFR exhibit smaller sequence conservation scores than ATP binding sites which indicates that more sequence variability can be tolerated for folding initiation sites. Again, protein function depends on particular amino acid side chains [83], whereas protein structure and secondary structure element formation is mainly the consequence of the hydrophobicity of amino acids [94,95]. ATP binding sites exhibit lower EFR prediction scores compared to the average in the protozyme region which captures their tendency to occur in exposed, unordered coil regions as observed in the previously reported findings.

**Fig 7.**
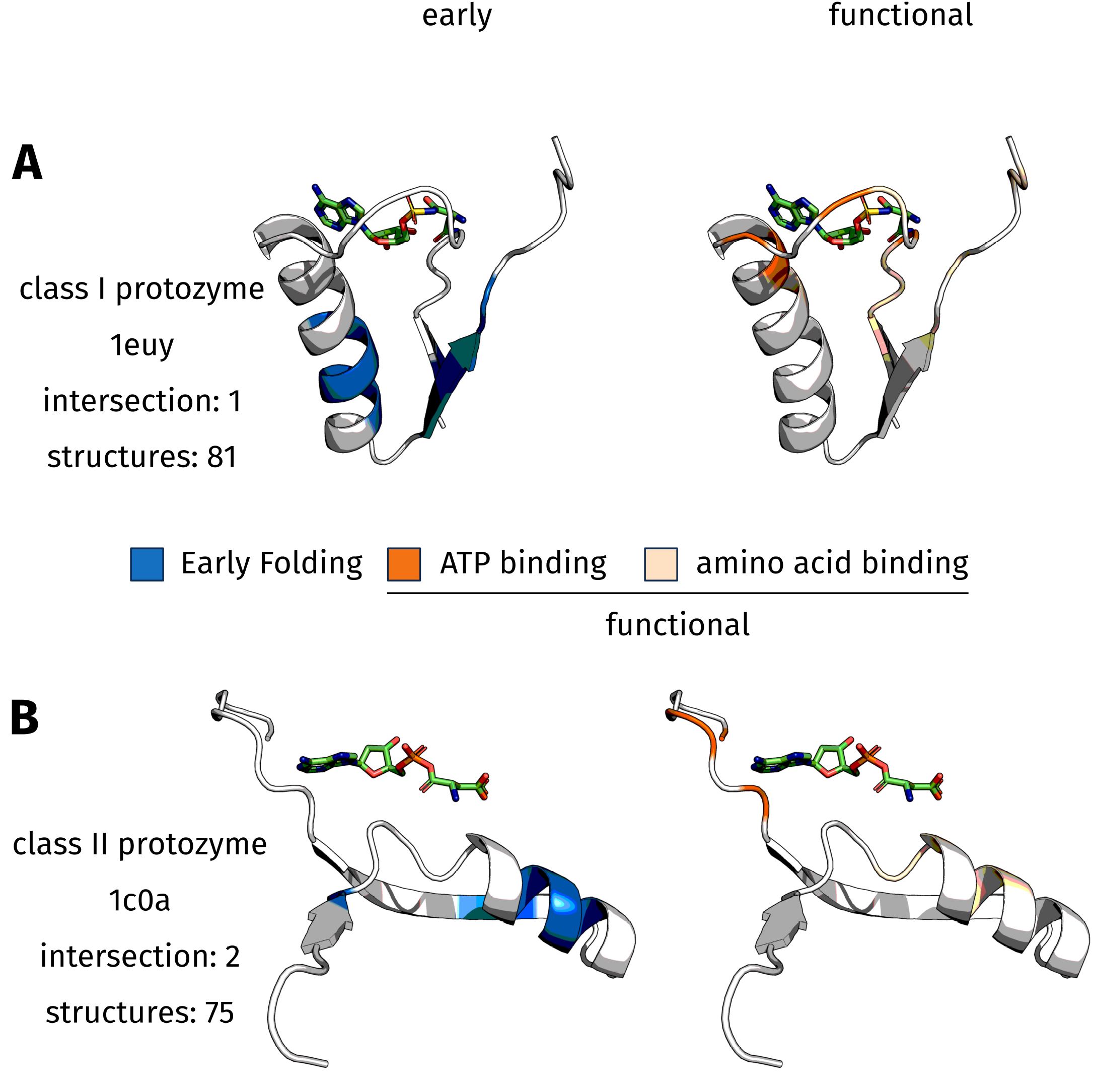
Hypothesized protozyme regions of both aaRS classes. The protozymes [80,81,104] (in cartoon style) and the respective aminoacyl-AMP ligand (in sticks style) are depicted. This captures the state after the first reaction after ATP and amino acid have been covalently bound. The ATP part is oriented to the left, whereas the amino acid is located on the right. Residues predicted to be Early Folding [9] are colored blue, whereas functional residues [104] are rendered in orange. ATP interaction sites are depicted in dark orange, residue positions observed to interact with the amino acid in any aaRS structure are rendered in light orange (see S6 Fig for a schematic depiction). In the rare cases that residues are both EFR and functional, they bind the amino acid part of the ligand in two specific aaRS implementations. (**A**) The class I protozyme is represented by truncated PDB:1euy_A. The respective EFR as located in the center of the ordered secondary structure elements. In contrast, functional ligand binding sites are located in the upper part of each subfigure. They are primarily located in unordered coil regions. (**B**) The class II protozyme, represented by truncated PDB:1c0a_A, shows similar tendencies.

ATP refers to the number of ATP binding sites and aa refers to the number of positions realizing amino acid recognition in any aaRS implementation. The intersection of functional residues involved in ATP and amino acid binding is given. The shift in probability to the expected intersection is stated. A perfect separation of EFR and functional residues in the sense of ATP binding positions can be observed. Also, positions relevant for amino acid specificity are remarkably well separated from EFR most of the time. The overlap is present in the amino acid recognition sites in two implementations respectively: TrpRS and TyrRS in class I and AspRS and PylRS in class II.

In class I (visualized by truncated PDB:1euy_A), position 311 is the only residue which is both EFR and functional (Table 2). This position is only functional in TrpRS and TyrRS where it realizes binding of the respective amino acid. Both tryptophan and tyrosine are large, aromatic amino acids and it is hypothesized that they were added late to the genetic code [103]. This implies that these EFR became functional late during the evolution of aaRS. The clear separation with respect to ATP recognition implies that the unifying aspect of all aaRS is binding of the ATP ligand and catalysis at the respective *α*-phosphate [104]. At first protozymes where required to bind ATP and later the amino acid binding sites improved in specificity, allowing them to discriminate between amino acids more reliably. Position 274 corresponds to the N-terminal residue of the Backbone Brackets structure motif. Close to this position various amino acid binding sites can be observed in other class I aaRS, while EFR are further away (S4 File). Despite being functionally relevant, the sequence conservation of position 274 amounts to 3 and is relatively small. This residue has been shown to realize ATP binding by backbone hydrogen bonds which can be virtually realized by all amino acids [104]. Thus, change can be compensated at this position as along as the backbone atoms can still bind the ATP ligand. Furthermore, this position interacts with the *α*-phosphate position of the ligand to which the aaRS attaches the proper amino acid [104]. Therefore, it is intuitive that many positions involved in amino acid recognition are located at neighbored sequence positions. In class I, 15 of 16 EFR positions in the MSA relate to well-mapped positions (i.e. present in the majority of aligned sequences). Moreover, the LFR position 284 features a remarkably high sequence conservation of 10. This position is part of the HIGH sequence motif which relates to ATP binding and the stabilization of the transition state [102]. In most class I aaRS, the HIGH motif is located at the C-terminal end of a *α*-helix. Despite this defined secondary structure, the HIGH motif is predicted to consist of LFR. EFR are located in the center of the helix: sites, with a high drive to fold, will initiate the formation of helices and then extend them until the sequence composition hinders any further extension [7,8].

**Table 2.** Comparison of folding characteristics and functional relevance for aaRS classes.

| class | early | ATP | aa | ATP int. | aa int. | ATP shift | aa shift |
| --- | --- | --- | --- | --- | --- | --- | --- |
| class I | 16 | 4 | 13 | 0 | 1 | -0.95% | -1.87% |
| class II | 10 | 4 | 8 | 0 | 2 | -0.82% | 1.22% |
|  | 26 | 8 | 21 | 0 | 3 | -0.90% | -0.39% |

In class II, positions 665 and 666 are both functional and predicted to be EFR (Table 2). Again, these positions are not functional in most class II aaRS. Only in AspRS and PylRS they are observed to bind the amino acid part of the ligand. In agreement with the observation for aaRS class I and the acyl-coenzyme A binding protein, asparagine and pyrrolysine are relatively large ligands which may require EFR to participate in protein function. 9 of 10 EFR positions are well-mapped in class II. For both classes, functional positions are well-mapped as well in the majority of observations. For position 698 of class II a sequence conservation score of 11 is observed. This position is the N-terminal residue of the Arginine Tweezers structure motif [104] which has been demonstrated to depend on the conservation of this particular amino acid for ATP binding via salt bridges and *π*-cation interactions. Similar to class I, ATP binding positions can be found accumulated together at sequence level without any EFR between them (S5 File).

The findings related to the protozymes of aaRS substantiate that the most important aspect of a protein during evolution is function [87] and not retaining a particular protein fold [88]. Functional residues (i.e. ATP binding sites consistently shared by all aaRS implementations) exhibit a higher sequence conservation than EFR. The separation of EFR and functional residues is perfect when amino acid binding positions are ignored which are only relevant in a small number of implementations. Even when these amino acid binding positions were considered to be functional in all implementations, the intersection is remarkably small for class I. In this diverse superfamily, EFR are located consistently in the same columns of the MSA which agrees with the observation that this characteristic depends on the composition of larger sequence fragments [9] and relatively insensitive to inserts. Furthermore, it is shown that the position of folding initiation sites is preserved over the course of evolution even when the corresponding sequence conservation is small. Folding initiation sites occur in the center of secondary structure elements, independent of aaRS class.

## Conclusion

A dataset of Early Folding Residues for the protein folding process was studied. They are highly connected nodes in residue graphs and were observed to be located in energetically favorable conformations as pointed out by the approach of Energy Profiling [28,29]. These structurally relevant residues have distinct properties e.g. regarding the number of hydrophobic interactions compared to functional residues.

Future hydrogen-deuterium exchange data can substantiate the presented trends regarding the nature of EFR. Potentially, the arsenal of experimental techniques to study the folding process of proteins will expand and become more refined and standardized, so that the underlying dataset of studies like this one will become more robust. Early Folding Residues are an excellent tool to gain insights into the folding process with spatial and temporal resolution. Future studies may link them to characteristics at sequence level to understand the sequence composition which causes particular regions of a protein to initiate the folding process. Features presented in this study were shown to be discriminative for Early Folding Residues. Classifiers for them based on sequence [9] or structure may annotate residues crucial for protein folding. Trained classifiers can also report as well as visualize the most discriminative features [106,107] which may further delineate EFR. This information is also invaluable for mutation studies, *ϕ*-value analysis, or protein design and can serve as basis for the prediction of mutation effects [99]. The same is true for the observed separation of structurally relevant and functional residues in proteins. Understanding these topological differences provides insights into the way they interact with the rest of the protein and to what degree they tolerate or compensate manipulation. For decades, scientists longed for a glimpse into the folding process [7–9] and the analyzed dataset [40] provides just that. The experimental signals of early folding events are still difficult to interpret and the analyzed dataset may not be generalizable for large proteins, but the made observations indicate that Early Folding Residues are also relevant for the stabilization of the native structure. In an evolutionary context as captured by the analysis of the aaRS dataset [104], the positions of Early Folding Residues are consistent among homologues which implies that folding initiation sites are preserved over the course of evolution even though their sequence conservation is relatively small compared to functional residues.

## Methods

### Dataset creation

Folding characteristics of residues were obtained from the Start2Fold database [40]. Therein, the authors adopted the definition of EFR from Li et al. [30] and presented a refined dataset which ignores possible back-unfolding and aggregation events [108]. The database covers all structural protein families present in CATH and SCOP [9]. However, the size of the deposited proteins [9,23] varies from 56 to 164 residues (S1 Table) which likely makes this resources only relevant for the folding of similarly small proteins.

This procedure resulted in a dataset for EFR characteristics encompassing 30 proteins and 3,377 residues - 482 of the EFR class and 2,895 of the LFR class. Due to the nature of the HDX experiments no data can be obtained for proline residues which feature no amide group susceptible to HDX [38], rendering them LFR in any case. Annotation of functional residues was performed using the SIFTS [109] and UniProt [63] resources. For 23 proteins an annotation of binding sites or regions existed, totaling in 2,807 residues - 130 classified as functional and 2,677 as non-functional. A detailed summary of the dataset is provided in S1 Table. Information used from the Start2Fold database can be found in S1 File. Residues annotated as functional are summarized in S2 File.

### Graph representation and analysis

Protein structures are commonly represented as graphs. This allows a scale-invariant characterization of the neighborhood relation of individual amino acids in the context of the whole protein [50].

In this study, amino acids constitute the nodes of a graph, whereas covalent bonds and residue contacts are represented as edges. Residues were considered in contact when their C*β* atoms were less than 8 Å apart - if no C*β* atom was present the C*α* position was used as fallback. Furthermore, contacts were labeled as either local (i.e. the separation in sequence is less than six) or long-range (i.e. sequence separation greater than five) [110]. This distinguishes contacts stabilizing secondary structure elements and those which represent contacts between secondary structure elements. The set of distinct neighborhoods of a node is defined as all adjacent nodes which do not share any local edge to any element of the set. Betweenness is defined the number of shortest paths on the graph passing through a specific node, normalized by the number of node pairs [49,111]. Closeness of a node is defined as the inverse of the average path length to any other node [48]. The clustering coefficient of a node is the number of edges between its *n_k_* adjacent nodes divided by the maximal number of edges between *n_k_* nodes: 0.5 · *n_k_* · (*n_k_* − 1) [49].

### Feature computation

Energy Profiles were calculated from structure and predicted from sequence according to the methodology used in the eQuant web server [28,29]. Energy Profiles represent a protein’s complex three-dimensional structure as one-dimensional vector of computed energies. Thereby, the surroundings of each residue are characterized by one energy value. Therefor, the frequencies of an amino acid to occur buried or exposed to the solvent were determined. Using the inverse Boltzmann law, the fraction of both states can be expressed as pseudo-energy. The energy of a residue can then be computed by summing up the corresponding pseudo-energies of all interacting residues. Residues were considered in contact, when the distance of their *Cβ* atoms was less than 8 Å [28]. RASA values were computed by the algorithm of Shrake and Rupley [112]. Buried residues are defined as those with RASA values less than 0.16 [76]. Non-covalent residue-residue contacts were detected by PLIP [113]. Secondary structure elements were annotated using DSSP [114]. For both ASA and secondary structure element annotation the BioJava [115,116] implementations were used. The loop fraction is defined as fraction of unordered secondary structure in a window of nine residues around the evaluated amino acid [64]. This yields a fraction, where high values are tied to regions of high disorder, whereas amino acids embedded in *α*-helices or *β*-sheets result in scores close to zero. The centroid distance of a residue is the spatial distance of its centroid to that of all atoms. The terminus distance is lower of the sequence separation to either terminus divided by the number of residues. Evolutionary information as well as couplings scores were computed using the EVfold web server [65,66]. The evolutionary information is based on the MSA of homologues automatically retrieved for the query sequence and expresses how conserved a column in this MSA is.

Data integration was performed by a Java library publicly available at https://github.com/JonStargaryen/jstructure.

### Statistical analysis

Dependence of distributions of real-valued variables was tested by the Mann-Whitney U test. Dependence of distributions of count variables was tested using the Dunn test with Bonferroni correction. * corresponds to significant *p*-values <0.05 for the Mann-Whitney U and *p*-values <0.025 for the Dunn test. The observed intersection between EFR and functional residues was expressed as probability and compared to the expected probability of a residue to share both the EFR and functional label based on their respective probabilities to occur individually.

### Creation of the aminoacyl-tRNA synthetase dataset

An evolutionary trajectory of highly diverse proteins can be found in aaRS. A detailed description of the methodology can be found in a previous study [104]. 972 aaRS structures from the PDB were analyzed. Within each class, sequences were clustered together when their sequence identity was above 95%. For clusters of highly similar sequences, a representative was determined. All representatives within a class were aligned by the T-Coffee expresso pipeline [105]. Thereby, all structures were renumbered within each class and allows to directly compare structures: e.g. the first residue of the Backbone Brackets motif is at renumbered position 274 and can by found by that residue number in all other class I structures despite the high sequence variability. From these renumbered protein chains, the corresponding sequence was extracted and used as input for the EFoldMine algorithm [9] which predicts the probability of residues being EFR. This was necessary because no experimentally derived folding characteristics are available for aaRS proteins. Predicted scores exceeding 0.163 where considered EFR; this value has been shown to optimally separate EFR and LFR [9]. The annotation of functional residues was derived from a curated annotation of ligand binding sites [104]. For ligand binding, ATP binding sites and amino acid binding sites were distinguished as detected by PLIP [113]. Protozyme regions were extracted from PDB:1euy_A and PDB:1c0a_A to represent aaRS class I and II. This selection was for visualization purpose only and focused on structures which ligands in aminoacyl-AMP ligands. Selected residue numbers of the protozymes are 255-336 and 648-718, respectively [104]. In contrast to the evolutionary information scores [65,66], the sequence conservation in aaRS sequences was computed by Jalview [117,118] using only sequences which were used as input of the MSA. Positions composed of sets of amino acids with similar characteristics result in high values.

## Acknowledgments

The authors thank Florian Kaiser, Christoph Leberecht, Sebastian Salentin, and Alexander Eisold for scientific discussions and/or proofreading the manuscript.

## Supporting information

**S1 Fig. Amino acid frequencies.** The gray bars correspond to the amino acid frequencies in the dataset [23]. The blue bars depict the frequency of a particular amino acid to be an EFR. Hydrophobic amino acids show an increased tendency to be EFR.

**S2 Fig. Fine-grained distribution of computed energies.** A more fine-grained version of Fig 3. Additionally, the overall distribution of computed energies of an amino acid is depicted in gray.

**S3 Fig. Correlation matrix of computed features.** Depicts correlations of analyzed correlation. The bigger the circle, the higher the association of both variables. Blue refers to positive correlation, whereas red represents a negative correlation.

**S4 Fig. Network descriptors.** Depiction of the used network descriptors: betweenness, closeness, clustering coefficient, and distinct neighborhood count.

**S5 Fig. Dataset entries where EFR and functional residues are well-separated. EFR are rendered in blue, functional residues are rendered in orange.** (**A**) In the case of fibroblast growth factor (PDB:1rg8_A) EFR are distributed throughout the sequence are accumulated in the core of the protein. In contrast, functional residues primarily occur in a N-terminal heparin-binding region. (**B**) Similar tendencies are present in Villin-1 (PDB:2vil_A). This time the functional binding site is located near the C-terminus and contains no EFR.

**S6 Fig. Binding site of aaRS enzymes.** ATP binding regions are depicted in dark orange, whereas amino acid specific positions are rendered in light orange. Figure adapted from [104].

**S1 Table. EFR dataset summary.** Summarizes identifiers [23] of each entry as well as the number of residues in the corresponding protein chain, the number of EFR and functional residues as well as the cardinality of the intersection of both sets. In order to assess the relevance of the observed intersection it was compared to the expected intersection. Proteins not containing any functional residues according to UniProt [63] are marked with dashes.

**S2 Table. Statistical characterization of EFR.** For each presented feature the mean (*μ*) and standard deviation (*σ*) of both the EFR and LFR category is reported. *P*_buried_ refers to the *p*-value of the test on residues buried according their RASA value, this was done because EFR have a tendency to be located in the core of a protein and without filtering all differences are significant. Features and employed tests are described in the Methods section.

**S3 Table. Comparison of EFR and functional residues.** For each presented feature the distribution of values is compared between functional and non-functional residues as well as EFR and functional residues. The corresponding *p*-values and significance level are stated for buried residues. Mean values are shown for EFR (*μ*_early_) and functional residues (*μ*_func_). Features and employed tests are described in the Methods section.

**S4 Table. Summary of the aaRS dataset.** Sequence conservation [117,118] and EFoldMine [9] predictions for the aaRS protozyme regions [80,81,104] are presented. Encompassed are the average values for all residues, residues in the protozyme region, for positions predicted to be EFR, functional residues, ATP binding residues, and amino acid binding sites.

**S1 File. Start2Fold dataset as JSON file.** Machine-readable JSON version of the dataset. Provides protein name, Start2Fold identifier, PDB identifier, UniProt identifier, number of EFR, range of residues numbers, and the secondary structure element composition for each dataset entry.

**S2 File. Start2Fold dataset as table.** Summary table of all protein chains used for the analysis. Provides Start2Fold identifier, PDB identifier, evaluated experiments, number of EFR, UniProt identifier, and identifiers of functional residues derived from UniProt.

**S3 File. Table of computed features for the Start2Fold dataset.** Contains for all residues the set of computed features as well as the annotation of Early Folding and functional residues.

**S4 File. Detailed description of aaRS class I structures.** For each renumbered position, it is stated whether it is functional [104] or an EFR. Furthermore given are the sequence conservation [117,118], the number of backing sequences [104], and the average EFoldMine score [9].

**S5 File. Detailed description of aaRS class II structures.** For each renumbered position, it is stated whether it is functional [104] or an EFR. Furthermore given are the sequence conservation [117,118], the number of backing sequences [104], and the average EFoldMine score [9].

